# A Vibrating Ingestible BioElectronic Stimulator Modulates Gastric Stretch Receptors for Illusory Satiety

**DOI:** 10.1101/2023.07.17.549257

**Authors:** Shriya S. Srinivasan, Amro Alshareef, Alexandria Hwang, Ceara Bryne, Johannes Kuosmann, Keiko Ishida, Joshua Jenkins, Sabrina Liu, Wiam Abdalla Mohammed Madani, Alison M Hayward, Niora Fabian, Giovanni Traverso

**Affiliations:** Department of Mechanical Engineering, Massachusetts Institute of Technology, Cambridge, MA 02139, USA; Division of Gastroenterology, Hepatology and Endoscopy, Brigham and Women’s Hospital, Harvard Medical School, Boston, MA 02115, USA; David H. Koch Institute for Integrative Cancer Research, Massachusetts Institute of Technology, Cambridge, MA 02139; Society of Fellows, Harvard University; Division of Comparative Medicine, Massachusetts Institute of Technology, Cambridge, MA 02139, USA

**Keywords:** vibrational stimulation, vagal activation, obesity, diabetes, gastrointestinal neurostimulation, neural interfaces

## Abstract

Effective therapies for obesity either require invasive surgical or endoscopic interventions or high patient adherence, making it challenging for the nearly 42% of American adults who suffer from obesity to effectively manage their disease. Gastric mechanoreceptors sense distension of the stomach and perform volume-dependent vagal signaling to initiate the gastric phase and influence satiety. In this study, we developed a new luminal stimulation modality to specifically activate these gastric stretch receptors to elicit a vagal afferent response commensurate with mechanical distension. Here we developed the Vibrating Ingestible BioElectronic Stimulator (VIBES) pill - an ingestible device that performs luminal vibratory stimulation to activate mechanoreceptors and stroke mucosal receptors, which induces serotonin release as well as yields a hormonal metabolic response commensurate with a fed state. We evaluated VIBES across 108 meals in swine which consistently led to diminished food intake (∼40%, p< 0.0001) and minimized the weight gain rate (p< 0.03) as compared to untreated controls. Application of mechanoreceptor biology could transform our capacity to help patients suffering from nutritional disorders.

## Introduction

The obesity epidemic, affecting nearly 42% of U.S. adults*(1, 2)*, increasingly strains healthcare resources by increasing the incidence of comorbidities such as diabetes, hypertension, cancer and heart disease*(3, 4)*. Given the difficulty of modifying behaviors and limitations in weight loss associated with pharmacologic therapies, there remains a pressing need for new methods that efficaciously decrease weight gain. While bariatric surgeries have demonstrated efficacy, and have evolved as minimally invasive laparoscopic procedures (Roux-en-Y and laparoscopic banding), they require significant pre- and post-surgical lifestyle modifications and remain too costly (7,400 USD and 34,000 USD) for the global populations requiring treatment (5–7).

Vagal nerve signaling plays a critical role in satiation through a negative feedback loop in which anorexigenic neurometabolic secretions are released in response to food intake*(8)*. Distension of the stomach by food contents is transduced by intraganglionic laminar endings (IGLEs), the most-prevalent type of vagal afferents innervating the gastric musculature, which sense contraction and distension*(9, 10)*. These stretch mechanoreceptors produce short-acting vagal afferent signals and increase neuronal activity in the nucleus of the solitary tract (NTS)*(11)* where vagal afferents terminate and interact with reward, energy homeostasis, hunger, and mood circuitry*(9, 12)*. In turn the NTS triggers metabolic and neural anorexigenic signaling*(13–15)* to yield feelings of hunger or fullness and alter food intake*(16)*. Since this mechanism is primarily volume-dependent*(17)*, as opposed to composition-dependent, (carbohydrates, proteins, fats, or saline*(18)*), methods to manipulate gastric volume-- intragastric balloons (IGB)--were developed as an easy-to-deploy tool*(19)* to minimize weight gain.

IGBs are designed to induce stomach distension to induce early satiety. Although they enable short-term weight loss during the adaptation phase, IGBs fail to promote sustained changes in hunger or eating behavior after 10-12 weeks*(20)* nor do they demonstrate superior outcomes compared to pharmacologic or surgical therapy*(19)*. Neural adaptation to the chronic distension (as opposed to periodic distension that results from eating), as well as placement, removal, perforation and obstruction complications pose challenges for the long term efficacy and safety of IGBs*(21, 22)*. Following numerous deaths in patients with IGBs since 2016, the FDA has issued warnings and some companies have recalled their IGB products*(23)*.

Intervening more proximally, vagotomies and electrical vagus nerve stimulation (VNS), which are more localized interventions, have demonstrated preclinically to be associated with decreased weight gain, food intake, and sweet cravings, and increased satiation and energy expenditure*(12)*. When clinically implemented for depression and epilepsy, VNS has shown a decrease in weight gain, reduced sweet cravings and increased energy expenditure. However, non-gastrointestinal (GI) side effects resulting from nonspecific axonal targeting at the cervical level of the vagus and metabolic compensatory mechanisms have prevented widespread clinical implementation for obesity*(24)*. Vagatomies have also demonstrated significant benefit*(25)*, although the mechanism and side effects are unclear and requires an invasive surgical procedure*(26)*. Fundamentally, with our current technology and understanding of neural signaling, VNS systems cannot specifically target the relevant axons nor perform patterned stimulation to recapitulate the complex physiological signaling underlying satiety.

Considering the central role of gastric mechanotransducers in vagal neurometabolic satiety signaling*(27)*, a mechanism and devices capable of selective mechanoreceptor activation would pose significant value. Seminal experiments in stretch-sensitive spindle fibers in skeletal muscle have shown that vibration elicits illusory distension*(28–30)*. Following a parallel mechanism, we devised a vibratory stimulation modality to selectively activate gastric stretch receptors and characterize their response in this proof-of-concept study. We hypothesized that optimized intraluminal vibration of the gastric smooth muscle would induce illusory distension of the stomach, generating vagal afferents signals and a metabolic response commensurate to those elicited from mechanical distension in a fed state (Figure 1a). We then developed the Vibrating Ingestible BioElectronic Stimulator (VIBES) pill, a safe, easy-to-use, ingestible device that can perform temporary, targeted, intraluminal mechanical stimulation prior to meals to achieve early satiety. In an awake and freely moving swine, we hypothesized that the VIBES would reduce food intake and minimize the rate of weight gain as compared to untreated controls.

**Figure 1.**
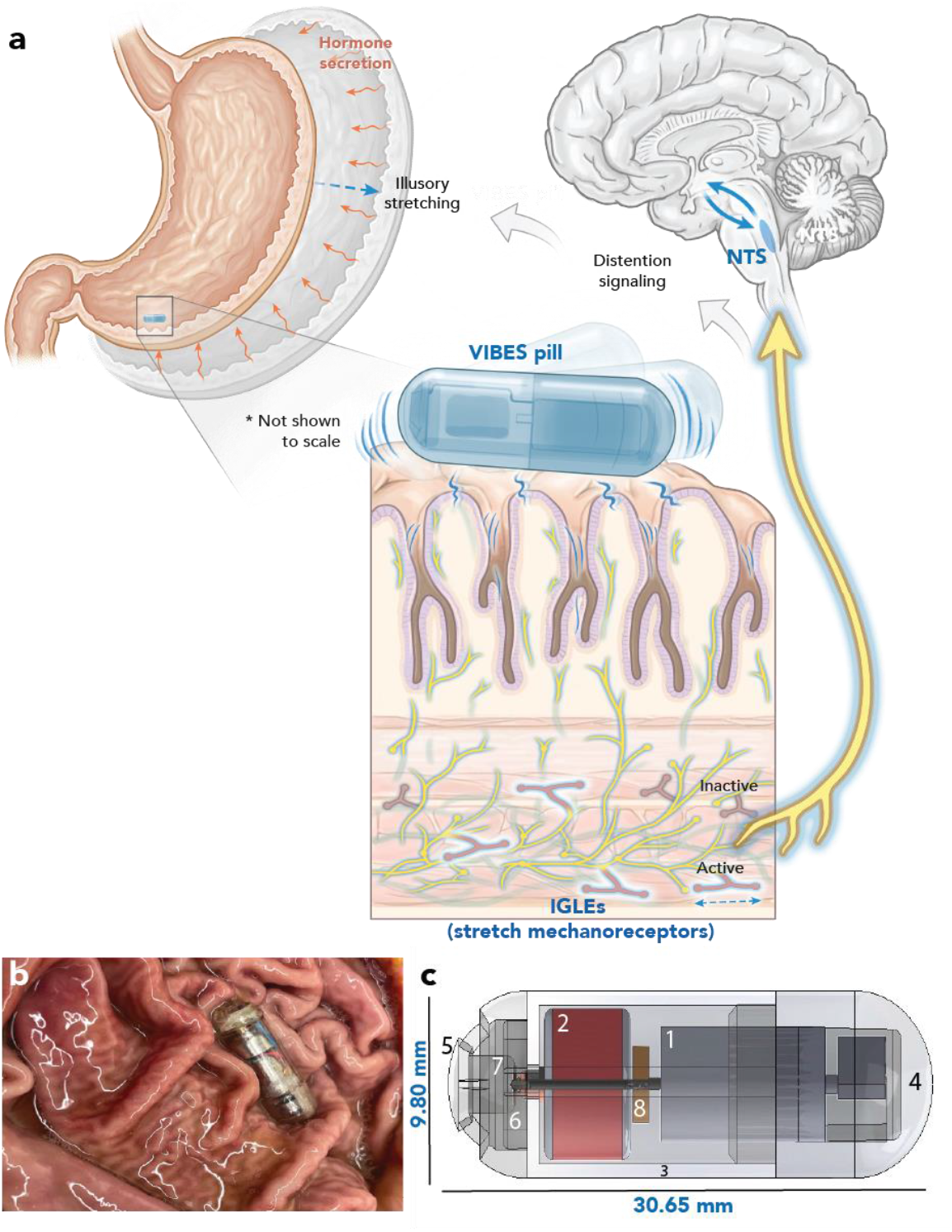
VIBES Concept and Mechanism. **a)** The VIBES makes contact with the gastric lining and activates following contact with gastric fluid. Vibrations activate IGLEs in the celiac plexus, signaling distension to the NTS, which interacts with hunger circuitry to signal illusory distension. b) VIBES sits amongst gastric rugae in a swine stomach and strokes the mucosa as it performs stimulation. C) The VIBES pill consists of a 1) offset motor 2) silver oxide battery 3) central body 4) motor cap 5) pill cap 6) pogo pin 7) gelatinous membrane and 8) resistor. Original illustration by Virginia Fulford.

## Results

### Design and Characterization of the VIBES pill

The VIBES was designed to be orally ingested, to sustain contact with the gastric lining, to activate upon submersion in gastric fluid, to vibrate with amplitudes sufficient to stimulate gastric IGLEs for a set time period, and to pass safely through the GI tract. Its triple-zero capsule houses a gelatinous membrane which dissolves in 4.3 +/- 1.2 minutes following immersion in gastric fluid, releasing a spring-loaded pogo pin that completes the circuit to activate the vibrating motor. Pill ingestion and travel through the esophagus is known to take a maximum of 60 seconds, ensuring that the pill will reach the stomach prior to activation*(31)*. A motor with an offset shaft is positioned within a custom housing enabling displacement amplitudes of 2-4mm when powered with a 1.55V 80 mAh silver oxide battery.

Duration testing was performed by immersing the VIBES pill in gastric fluid on a soft substrate. The pill actively vibrated for an average of 38.3 ± 1.83 minutes for the VIBES pill (n=5). Since meals are generally consumed in a 20-to 30-minute window and gastric contents undergo primary mixing in approximately an hour, this time range was determined to be acceptable. To ensure that the system did not degrade or have material vulnerabilities, a chemical resistance test (described in Methods) was performed by immersing the VIBES pill in simulated gastric fluid (pH = 1.2) for 24 hours and simulated intestinal fluid for 10 days, both at 37°C. No macro or microscale changes were observed following immersion and pills were able to be successfully activated after the incubation period (Supplemental Figure 1). This ensures that the VIBES pill would not damage the tract even if it were to reside in the stomach for a full day or in the intestines for over a week, much longer than the expected or measured residence times with normal motility. Thermal testing was performed to assess any potential heating risks on the surrounding tissue environment. 30 minutes of operation at various frequencies yielded less than a 0.5°C change in the surrounding fluid, ensuring no thermal risks for the mucosal layer during operation (Supplemental Figure 2).

A swine model (50-to 80-kg Yorkshire pigs ranging between 4 and 6 months of age) was utilized to study the VIBES performance as its gastric anatomy is similar to that of humans. Further it has been widely used in the evaluation of biomedical gastrointestinal devices*(32)*. Localization of the VIBES pill was characterized through endoscopic observation in n = 10 swine. VIBES was endoscopically deployed into the gastric cavity and the final positioning was observed through the video channel over the course of 30 minutes. The VIBES pill exerts 0.04214 N of downward force and has a density of 2.019g/cm^3^. In all 10 trials, the pill sank through gastric contents (densities 0.011 – 1.158 g/cm^3^) and achieved stable contact with the lining (Supplemental Note 1). Localization in the gastric antrum and cardia were common and dependent on the positioning of the animal. In several cases, over the course of 30 minutes of stimulation, the pill was seen to migrate along the lining. During the 30-minute period, trained endoscopists monitored the density of the ruggae in the empty stomach to estimate changes to the gastric cavity volume. In all trials, no differences in the stomach volume were visualized.

In all cases, the VIBES was not emptied from the stomach by phasic inter-digestive migrating motor complex (MMC) for at least 30 minutes following administration. Thus, if VIBES is taken prior to meals to start the distension response, it will remain present to interact with the gastric lining during intake and the primary mixing phases of digestion.

### Stretch-receptor Activation to Signal Gastric Distention

To characterize the neural signaling patterns of the stomach in response to mechanical distension and vibrational stimulation (n = 4 animals), fine-wire electrophysiological recordings of up to 24 branches of the celiac vagus nerve, which innervate the gastric cardia, were performed following laparotomy. The celiac vagus was selected to isolate signals arising from the stomach and mitigate off target signals from other organs. Baseline recordings demonstrated small amplitude, spontaneous spikes along with slow wave related spiking. To mimic the distension created by the intake of food, the stomach was inflated using the endoscope to 30%, 60%, and 90% of its maximal volume and held for 180 seconds (Supplemental Figure 3). Five minutes of rest were provided between each insufflation state to allow neural activity to return to baseline. Spiking was observed in a subset of 6-10 channels with onset occurring 10-12 seconds following the beginning of inflation (Figure 2a, 2b); these channels were designated as channels corresponding to axons innervating low-threshold stretch-sensitive IGLEs, consistent with prior studies*(10)*. Spiking amplitude was higher and more densely concentrated at the onset of stretch followed by a slow adaptation rate (levelling off of distention-based afferents as the stomach adapts), commensurate with prior reports of these afferents*(10)* (Figure 2b). Upon desufflation, spiking acutely decreased and terminated within 2-18 seconds. The rate of firing graded monotonically with distension which is commensurate with the known spiking patterns of IGLEs (Figure 2a) *(10)*. We modulated the distension by varying the volume of insufflation in the stomach

**Figure 2.**
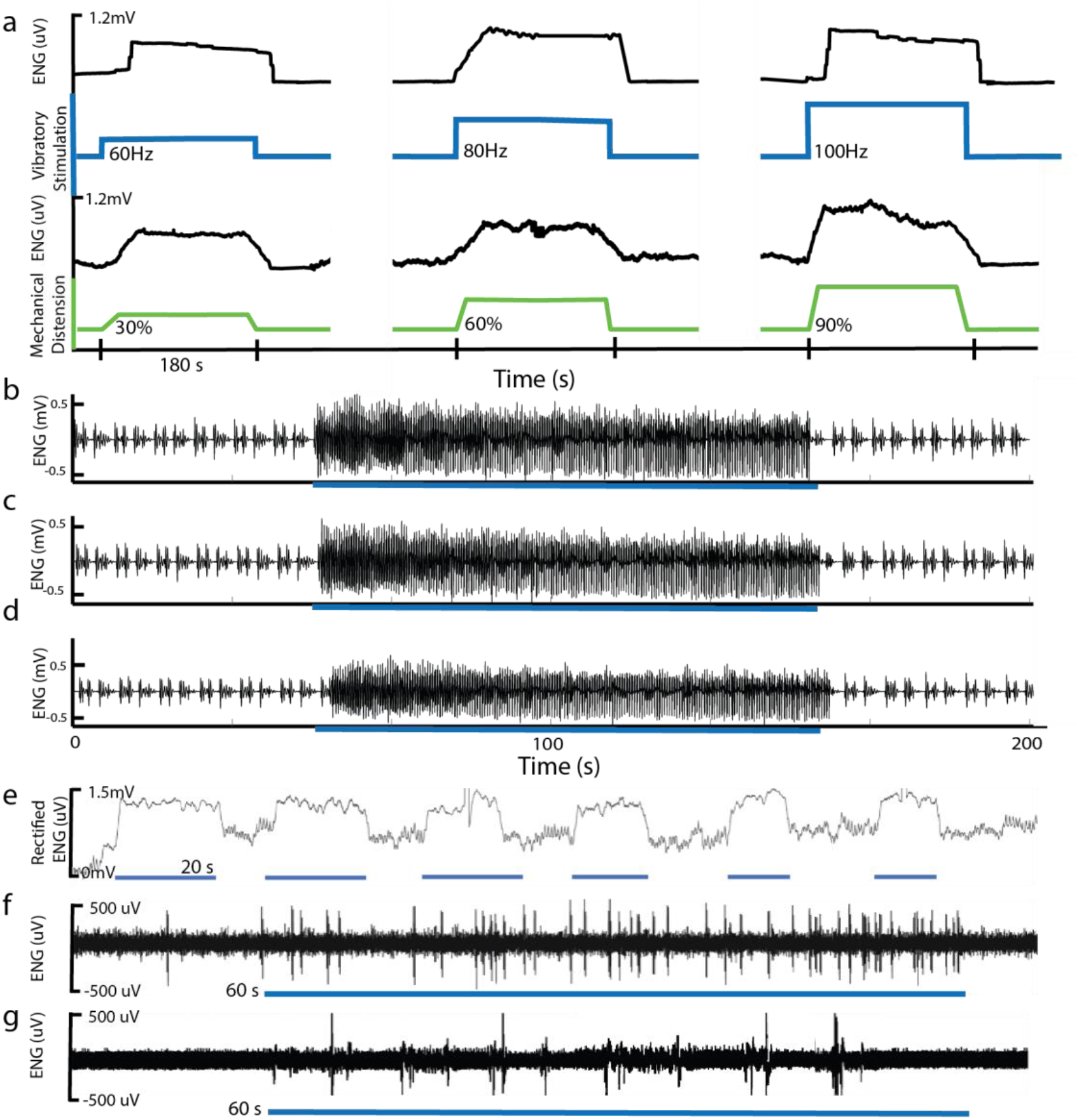
Gastric Afferent Electrophysiology of Stretch-Sensitive Mechanoreceptors. a) The gastric cavity (green) was distended 30%, 60%, and 90% of the gastric volume using insufflation. The corresponding electroneurographic (ENG) vagal response was recorded from celiac vagal branches demonstrating spiking in response to distension. Vibratory stimulation at 60, 80, and 100 Hz (blue) resulted in similar, monotonically graded ENG responses (black). ENG signals have been rectified for visualization. b,c) ENG signal from a stretch sensitive afferent fiber responding to (b) mechanical inflation and (c) a VIBES pill within the stomach and (d) a VIBES pill after bilateral vagotomy. Heart rate and respiratory artifact are present at baseline e) Periodic VIBES vibration resulted in repeatable induction of the stretch response. Afferent response to mucosal stroking of the gastric lumen by an (f) endoscopic fiber and (g) VIBES pill rotation against the mucosal surface. Blue bars underlying electrophysiological traces indicate when stimulation was active.

To determine if vibratory stimulation could activate these IGLEs to produce a similar response, neural activity on the channels that were active for stretch-sensitive IGLEs were monitored as vibration with frequencies between 24 and 500 Hz was applied using the VIBES to the gastric lumen (Supplemental Figure 4). Although the pill provides a contact surface area of approximately 298 mm^2^, the vibratory stimulation propagates through tissue, stimulating the gastric muscularis up to 30 cm away.

In all trials, spiking was observed at frequencies between 64 - 100 Hz in a similar pattern to those observed during mechanical distention (Figure 2c). At these frequencies, the amplitude of displacement of the pill was greater than 1mm. No activation was observed for displacements under 1mm, or for frequencies below 35 *Hz* or above 150 Hz. Onset of activation occurred on average within 2.5-3.2 seconds. The change in signal amplitude demonstrated slow adaptation, similar to mechanically-activated afferents (Supplemental Figure 5). Bilateral vagotomy was then performed and the above results were replicated to eliminate potential confounding between efferent signals and feedback loops. Raw spiking patterns from (Fig 2b) mechanical insufflation (Fig 2b), VIBES (Fig 2c), VIBES after bilateral vagotomy (Fig 2d) demonstrate a high degree of similarity in shape, pattern, and frequency. Furthermore, spiking was able to be elicited repeatedly (Figure 2e), without significant reduction in the onset or plateau amplitude (p < 0.05, n = 10). The frequency spectra of the average signal generated in the control and VIBES groups were compared to assess the similarity of the afferent evoked response (Supplemental Figure 4). The dominant peak was identical in both groups at 2904 Hz with similar amplitudes. Further, the 5 greatest peaks occurred at the same frequencies in both groups. Cross-spectral coherence between the VIBES and control afferents was 0.0224, which is not statistically different at a p-value of 0.4426. A two-tailed t-test of the average signals was also insignificant at an alpha of 0.1. These results suggest that gastric luminal vibration induces afferent neural activation of IGLEs characteristic of spindle-type reception of distention.

### Gastric Mucosal Stroking

In addition to distension, mucosal stroking triggers gastric mechanoreceptors to stimulate gastric secretory activity. Gastric mucosal stroking was conducted via an endoscope using a thin filament that is known to elicit spiking for such receptors (Figure 2g). During VIBES treatment, in axons not activated by stretch, we observed such periodic bursting (Figure 2f). The surface geometry of the VIBES pill may have stroked the gastric mucosa as it rotated, resulting in short, periodic bursts from quickly adapting mucosal receptors*(33),(11)*. Mucosal stroking is known to release 5-HT or serotonin*(34–36)*, which acts on vagal 5-HT_3_ receptors that perform satiation signaling as well as enteric 5-HT_4_ receptors which regulate peristalsis, secretion, vasodilation and digestion through intrinsic central and peripheral reflexes. We assayed the luminal secretion of serotonin in response to VIBES using ex vivo tissue on a Franz cell apparatus*(37)*. 80 Hz resulted in the greatest increase in secretion levels as compared to the control condition of 0Hz (Supplemental Figure 6). Based on this data, and the inconvenient human audibility of the VIBES pill above 100Hz, 80 Hz was selected as the optimal operating frequency. Additionally, surface geometries were altered to enable greater stroking of the mucosa (Supplemental Figure 7), including studs and ridges to enhance surface contact. A spiraling design induced a significant increase in luminal serotonin release as compared to the straight surface and control (untreated) conditions.

### Metabolic Effect of VIBES

To characterize the downstream effects of VIBES through vagal afferent signaling, we profiled the hormonal secretions relevant to feeding behavior and satiety. Blood was sampled at 0, 15, 30, 45, 60, 90, and 150 minutes for animals receiving a VIBES pill (n=6, between 30-60^th^ minute marks) and a sham pill (n=6). The effect was ascertained using a two-tailed heteroscedastic t-test of the average value of the hormone after stimulation between the control and VIBES groups at an alpha of 0.05. After stimulation, the pill remained in the tract and was allowed to pass naturally. While the control animals exhibited the expected levels of these hormones in a fasted state, treatment with VIBES resulted in a significant reduction of ghrelin, the ‘hunger hormone’ (p < 0.01). Ghrelin decrease is normally a postprandial response but occurred here in *fasted* animals treated with VIBES. Furthermore, insulin levels increased significantly (p < 0.01) with an amplitude and rate commensurate with normal meal ingestion. Glucagon, responsible for maintaining blood glucose levels, increased until minute 60 and then decreased as time progressed. Levels of C-peptide, associated with the biosynthesis of insulin, GLP-1 which enhances insulin secretion, and Pyy, an appetite suppressant, increased significantly with stimulation (p < 0.01). Together, these trends suggest that by signaling distention artificially, VIBES can induce the gastric phase metabolic response. Glucose from a venous catheter was measured at 30-minute intervals before and after administration of VIBES. No animals demonstrated hypoglycemia during the course of the study.

### VIBES reduces food intake

To investigate the effect of VIBES on hunger and feeding behavior, the food intake of 4 swine was monitored for at least 24 meals with no treatment (PEG-control group) and when treated with VIBES tethered through a PEG tube VIBES group). Tethering was performed with a very flexible leash of 10-15cm, enabling it to excurse around in the stomach freely. Endoscopic observation the free and tethered VIBES demonstrated no significant different in contact, vibration, mechanical force transmission and/or stroking of the gastric lining. Stimulation was performed for 30 minutes prior to mealtimes at 7:30am and 3:30pm when the hopper would be filled with pellets. Additionally, a snack of 5 apples was provided between 11am-12pm. While animals were fed ad libitum, the proportion of food consumed in the first 30 minutes after provision was measured to be the intake for a given meal. 4 animals matched in size and age were also monitored as controls, to account for potential effects of the PEG tube (control group).

Prior studies have demonstrated that satiety is well-marked by the amount of food consumed in animals*(38)*.The average percentage and standard deviation of the meal consumed for the VIBES, PEG-control, and control group were 58.1 +/- 10.7 (n=108 meals), 84.1 +/- 4.5 (n = 100 meals) and 78.4 +/- 4.5 (n = 96 meals), respectively. Intake in the VIBES group was significantly lower than the PEG-control and control groups (p < 10^−20^, student’s two-tailed homoscedastic t-test, Figure 4a). There was no significant difference between the PEG-control and control groups (p>0.05), indicating the presence of a PEG tube itself did not confound the significant difference seen during VIBES treatment. The distributions in the intake of each group were compared using the Kolmogorov-Smirnov test to further characterize the effect of the VIBES intervention. Between the VIBES and control groups, the p-value of 6.0103e-07 indicates that the distributions are significantly different with a maximal vertical distance between the cumulative distribution functions (CDFs) of 0.4232. However, between the PEG-control and control groups, a p-value of .055 and maximal distances between CDFs of 0.2119 indicate that there is no significant difference between these distributions (Supplemental Figure 17). On a per-animal basis, VIBES treatment resulted in significant reductions in intake (p< 0.001 in all cases, student’s two-tailed homoscedastic t-test), averaging 31% of the usual intake. Energy consumed per meal by each animal treated with the VIBES was also significantly lower than the control groups (Figure 4c, p< 0.001 in all cases, student’s two-tailed homoscedastic t-test). No adaptations or trends in intake were observed over the 24-meal period. To assess latencies and potential long-lasting effects of the treatment, a cross-over study design was utilized in which swine were treated for 3 meals, untreated for the subsequent 3 and treated for the final 3 (Figure 4d). Intake sharply increased during the untreated window, suggesting that VIBES functions through temporal vagal activation, with little neural adaptation or long-term effect. We also analyzed the trend of intake over the course of the study for each subject using Pearson’s correlation coefficients. No animals in any group demonstrated a significant trend in intake pattern (alpha = 0.1). This suggests that there is no habituation or adaptation to the VIBES treatment. The weight gain rate during the VIBES treatment period was significantly lower than the control period (p<0.03, student’s two-tailed paired t-test, Figure 4e) and significantly lower than the control group (p<0.01, student’s two-tailed heteroscedastic t-test). Together, these data suggest that the VIBES pill significantly decreases food intake and slows the rate of weight gain in a large animal model.

A preliminary behavioral study was conducted using an image-based deep learning and statistical model trained on 12 continuous hours of labeled daytime data (7a - 8p). This was used to analyze 96 hours of unlabeled daytime video data from 4 pig subjects in either the treatment or control conditions. We analyzed the time spent within each of four behaviors, detailed in Table 1 in terms of occurrences, durations, and probabilities within the control and treated conditions. The average percentage of time spent in each behavior demonstrates a trend towards more time spent in and over the feeder in the treatment condition (Figure 4f). Although the length of feeding bouts are not correlated with satiety*(38, 39)*, they are perhaps a marker of adaptive behavior to the intervention. Further, stimulated animals slightly trended towards more inactivity. On an hourly basis, controls demonstrated a relatively consistent level of activity and feeding throughout the day, while both trended downward in the treatment group (Figure 4g, h). Such reductions in physical activity and foraging have been reported to reflect postprandial satiety in laboratory pigs*(39)*. As treated animals consumed less food, on average, we hypothesized that they may demonstrate more appetitive behaviors before or near mealtimes, corresponding with a higher drop in blood glucose levels*(39, 40)*. These behaviors include locomotion correlated with foraging, rooting and interacting with cage walls*(39, 40)* and are positively correlated with feed restriction*(41)*. We categorized these behaviors in the active category and determined probabilities of a given behavior on an hourly basis. We observed peaks in the probability of active behaviors near mealtimes, potentially indicative that treated animals having consumed less food in a prior meal were ‘hungrier’ for the next meal (Figure 4i,j). The probability of inactive behavior also trended downward in treated animals over the course of the day and could be related to compounding or prolonged postprandial satiety, which is correlated with stabilized insulin and glucose levels measured (Figure 3)*(40)*.

**Table 1.** Behavior Classification.

| General Category | Behavior Type | Behavior Description |
| --- | --- | --- |
| Inactive | Sleeping | No movement, not standing, eyes open |
|  | Resting | No movement, not standing, eyes closed |
| Feeding | Free feeding | Head is in food bin, chewing |
|  | Pedialyte | Pedialyte through cage doors while in-vivo team exchanges batteries / fixes bandages: head touching bottle |
| Drinking | Water Spigot | Head to water spigot |
| Active | Hanging Toy Ball | Head or body touching the hanging ball |
|  | Hanging gear | Head or body touching the gears |
|  | Free moving toy | Interaction with head / body; moving |
|  | Scratching / rooting woodchips | Pawing at or using their nose to move woodchips |
|  | Scratching or climbing on walls of cage | Body up and touching the walls of the cage |
|  | Interaction with researcher | Touching, nosing, being fed from researcher |

**Figure 3.**
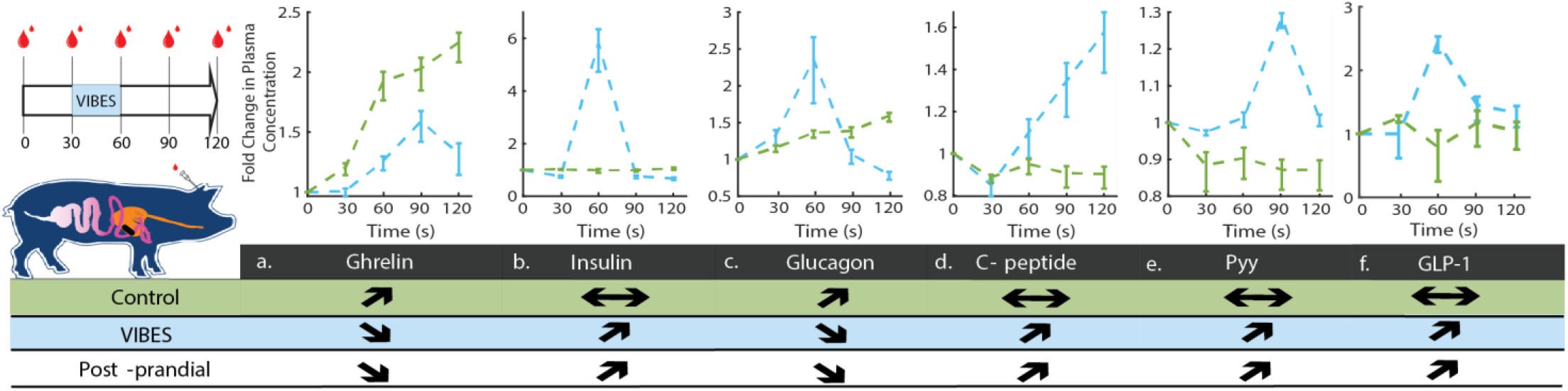
VIBES neuromodulation of gastric vagal afferents yields illusory metabolic satiety. a) Experimental schematic for blood sampling during VIBES stimulation between 30-60 minutes b-f) Response of hormones normalized to their baseline levels in animals with no stimulation (green) and VIBES (blue). Arrows indicate the observed trends for the Control and VIBES groups, and represent the known trend in animals and humans post-prandially.

**Figure 4.**
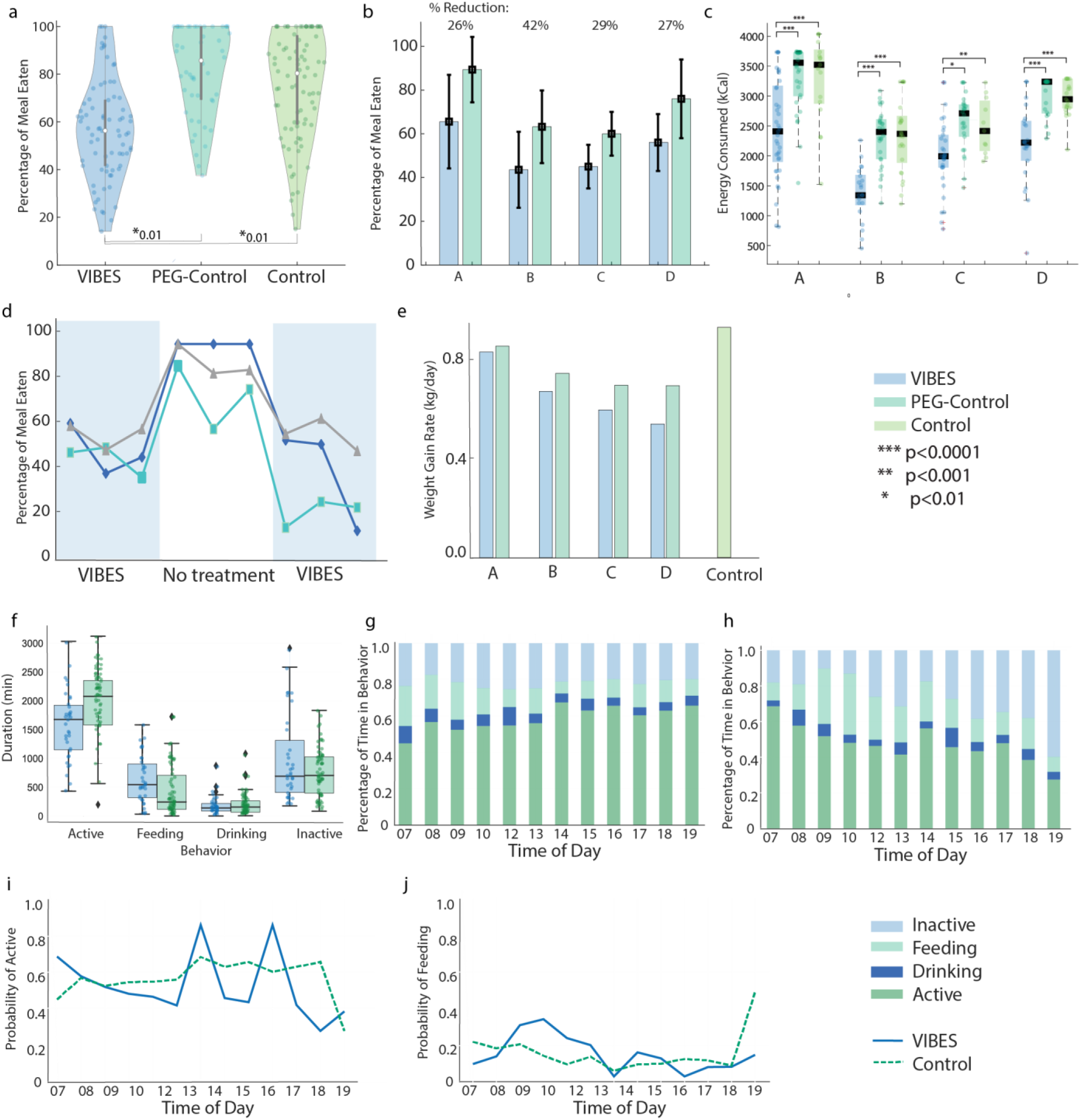
VIBES effect on feeding and weight gain. a) Percentage of meal consumed by swine in the VIBES, PEG-control, and control groups. b) Percentage of the meal consumed by animals A, B, C, and D over two weeks treated on VIBES and two weeks with no treatment (control). c) Energy consumed at each meal (dots) by animals A, B, C, and D over two weeks treated on VIBES, PEG-control or no treatment (control). Bars represent median and quartiles. d) Consumption of animals A (green), B (gray), and C (blue) in a cross-over study design comprising 3 meals with VIBES treatment, no treatment and VIBES, consecutively. e) Weight gain rate for each animal when treated with VIBES and no treatment. f) Duration of time that animals spent in each category of behavior. Percentage of time spent in each behavior for the g) control group and h) VIBES group. i) Probability of active behavior at each hour. j) Probability of feeding behavior at each hour.

### Safety and Biocompatibility

A blinded endoscopist assessed the lining of the stomach and surveyed for inflammation, degeneration, morphological changes, and other potential adverse effects from the pill in control (n=6) and VIBES animals (one administration of 30 minutes stimulation) (n=6) during endoscopy. No abnormalities were noted in either group. Even after two weeks of daily VIBES usage (twice a day), no abrasion, irritation, or inflammation was observed on endoscopic examination of the gastric cavity ([Supplemental video1](https://www.dropbox.com/s/0lfkw6k28qaosiv/Tour%20of%20the%20stomach.mp4?dl=0), [supplemental video 2](https://www.dropbox.com/s/yvrak5zln06llu9/StomachAfterVIBESbiocompatible.mkv?dl=0)). Further H&E staining on fixed sections of tissue following explant revealed no aberrant morphology, irritation or inflammation (Supplemental Figure 8).

To assess potential effects on motility and passage safety, animals were either fed a sham pill (n = 3 swine) or the VIBES pill (n = 3 swine) along with a set of small radio-opaque barium pellets. Radiography was performed every two days determined the period required for clearance of all barium pellets (Supplemental Figure 9). In animals with VIBES, swine passed all the pills in 4.3 days on average (range 4-5 days), while in control animals, passage required 8.3 days on average was required for sham pill passage (range 7 – 9 days) (Supplemental Table 1). Further, throughout the course of all trials in this study, veterinary staff monitored for changes in intake, health status, fecal output, lethargy, bloating and other behavioral signs of discomfort or disease daily. No adverse effects or discomfort was observed in the treated animals. No changes in stool quality or diarrhea were observed in any animals in the experimental or control groups. These data suggest that VIBES does not have any negative impact on motility. In all trials, animals were able to pass the pill without obstruction, perforation, or any signs of distress.

The pH of gastric fluid was monitored using an ex vivo Franz cell apparatus to assess whether the VIBES created any significant shifts in the gastric fluid. Following 20 minutes of stimulation in the experimental group, the pH of gastric fluid was not significantly different from that of the untreated control group using a student’s two-tailed heteroscedastic t-test (n=6 samples per group). The mean and standard deviation of the pH in the control and experimental groups was 1.265 +/- 0.03 and 1.260 +/- 0.029, respectively. As demonstrated by electrophysiology and the 5-HT response, the VIBES likely activates 5-HT_3_ receptors, which trigger critical gastrointestinal functions such as pancreatic secretion, meal termination, early satiety, and appetite regulation.

However, it is known that excessive activation of 5-HT_3_ results in nausea and vomiting *(42)*. Thus, all animals were monitored during treatment by 4-6 staff periodically during the day and through continuous daytime and nighttime video recordings 24 hours of the day. No signs of distress or emesis, or diarrhea were observed in any animal.

## Discussion

In this proof-of-concept study, we established a modality of luminal vibratory stimulation that activates gastric stretch receptors to signal distension and initiate the gastric phase. By optimizing the range of vibrational frequencies and including features in the VIBES pill that increased mucosal interactions, we not only generated vagal afferents signals relevant for indicating distention, but also induced a significant and consistent decrease in food intake in swine. Restriction of caloric intake during meals is a well-documented and sustainable mechanism to limit weight gain. We envision the VIBES pill being ingested on a relatively empty stomach 20-30 minutes prior to anticipated meals to trigger the desired sensation of satiety early in the meal. Shaping this luminal stimulation modality into a pill format presents several valuable advantages over its alternatives. As an ingestible device, no invasive implantation or surgery is required. Stimulation can be performed directly in the gastric cavity with a triggered activation, making the stimulation specific to the tissue of interest.

At scale, with injection molding techniques and mass manufacturing of electronics, the cost is expected to be in the cents to dollar range. Coupled with natural passage this makes the VIBES a consumable device, requiring no re-acquisition or recharging of the device. For certain patient populations, this enables temporary therapy, without the need for surgery. However, with improved power transfer and charging technologies, implantable or gastric-resident actuators could be developed to relegate the need for repeated oral administration for patients requiring chronic therapy.

The robust design of the VIBES overcomes practical limitations common to human oral consumption/administration. The 30Hz range of frequencies within which the stretch response is evoked is critical in enabling the VIBES’s function amidst varying gastric contents which may dampen the induced vibrations to varying degrees. The location of the VIBES pill is not controllable and likely to shift during a meal. This does not impede its function in the stomach as receptors in the antrum and cardia are predominantly stretch-sensitive*(10)*. Additionally, VIBES can consistently create a stretch response as gastric tension receptors are slowly adapting as compared to mucosal mechanoreceptors, which fire in short bursts as seen in our and prior studies*(11)*. The large animal study demonstrates that over a two-week period, and despite variations in the animal’s routine, sleeping patterns and activity, food intake was consistently lower when treated with VIBES. The response to the intervention also reinforces earlier studies highlighting the predominant influence of gastric distension in the satiety response.

Use of the VIBES resulted in no observable distress or negative side effects and normal passage in more than 20 trials in a large animal model, supporting the pre-clinical safety in a relevant animal model.

Weight loss studies are most commonly conducted in rodent models, given practical constraints of cost and resources, similarity in physiology, and ease of dietary-induced disease models*(43)*. Our choice of animal model was the swine, given the need for human-sized anatomy accommodating the geometric dimensions of the VIBES pill. However, weight loss is difficult to measure meaningfully in the young and growing swine species that are used for laboratory research*(44)*. Future studies will devise miniaturized VIBES devices apt for smaller animal model studies, be performed in a dog model which has a stomach geometry more similar to the human and utilize metrics like scintigraphy of liquid and solid meal, and gastric accommodation to further characterize the physiological effects. Future studies should delve deeper into potential correlations between the observed increases in inactive behaviors and mechanisms of postprandial somnolence. Additionally, the effect of the VIBES traversing through the small intestine on nutrient absorption, microbiome changes, and overall function of the GI tract should be thoroughly investigated. Mechanisms to externally control the activation could also be developed to increase safety and convenience.

Following further safety validations, clinical translation could facilitate a paradigm shift in potential therapeutic options for diseases such as obesity, polyphagia and Prader-Willi syndrome in which late onset of satiety yields excessive overeating and subsequent metabolic, cardiac, and endocrine comorbidities. Future studies should compare the efficacy and side effects of the VIBES pill to other FDA-approved drugs for weight control or emerging approaches like electrical stimulation*(45)*. The metabolic response triggered by VIBES could also be leveraged to treat diseases of insulin insufficiency or dysregulation such as diabetes. Its ability to increase the motility rate should be further studied. Optogenetic activation of the IGLEs could be another way in in which we could modulate the receptor physiology in the stomach.

Overall, this study lays the foundation for a new modality of vagal stimulation, acting through gastric mechanoreceptors, to induce an illusory sense of satiety, decrease food intake and limit the rate of weight gain, paving the way for a new treatment for obesity.

## Data and materials availability

All data associated with this study are presented in the manuscript or the Supplemental Materials.

## Acknowledgements

We are grateful for constructive feedback from Prof. R. Langer. Original artwork and Illustrations by Virginia E. Fulford, Alar Illustration. We thank Harrison Sun for his assistance with labeling data.

